# Convergent gliding, divergent ecology: Environmental drivers of gliding vertebrates in Southeast Asia

**DOI:** 10.64898/2026.04.30.721856

**Authors:** Kota Nojiri, Haruto Sugeno, Keito Inoshita

## Abstract

Gliding has evolved repeatedly across vertebrates and is often regarded as a classic example of convergent evolution associated with arboreal habitats. However, it remains unclear whether convergent locomotion corresponds to shared ecological responses across taxa. In this study, we investigated the distribution patterns and environmental drivers of gliding vertebrates in Southeast Asia using occurrence records and environmental variables representing climate and forest structure. We analyzed five major groups, including flying lemurs, flying squirrels, gliding lizards, gliding snakes, and gliding frogs, using presence-background logistic regression models. Across taxa, temperature seasonality showed consistently negative effects, while canopy height showed positive effects, indicating a shared association with climatically stable environments and well-developed vertical forest structure. In contrast, other environmental variables exhibited substantial taxon-specific variation. For example, elevation showed a strong negative effect only in gliding snakes, suggesting a tendency toward lowland habitats, whereas precipitation variables had limited explanatory power for gliding frogs. These results demonstrate that, despite the convergent evolution of gliding locomotion, ecological responses to environmental factors are not uniform across vertebrate taxa. Instead, species distributions are shaped by a combination of shared functional constraints and lineage-specific ecological traits. Our findings highlight the importance of vertical forest structure and suggest that habitat alteration affecting canopy structure may disproportionately impact certain taxa.

## Introduction

Gliding has evolved multiple times in the animal kingdom and is considered a classical example of convergent evolution (Rayner 1988; Dudley et al. 2007). A wide range of taxa, including insects, spiders, mollusks, fish, amphibians, reptiles, and mammals, exhibit gliding behavior, often as an adaptation to arboreal environments. In vertebrates, gliding has evolved independently in several lineages, including frogs, lizards, snakes, and mammals, typically in forested habitats where aerial movement between trees provides an efficient mode of locomotion (Rayner 1988; Dudley et al. 2007). The repeated evolution of gliding across distantly related taxa suggests that this locomotion is closely associated with specific ecological conditions, particularly those found in structurally complex arboreal habitats.

Southeast Asia represents a global hotspot of gliding vertebrate diversity and harbors a disproportionately high number of gliding species across multiple taxonomic groups (Emmons and Gentry 1983; Dudley and DeVries 1990; Wagner et al. 2023). This exceptional diversity has been linked to the unique ecological and evolutionary history of Asian tropical forests, particularly the dominance of dipterocarp trees and the structural complexity of the forests (Corlett 2007; Heinicke et al. 2012; Chaitanya et al. 2023). In contrast to other tropical regions, Asian forests are characterized by distinctive canopy architectures and ecological dynamics, which have been proposed to favor the evolution and persistence of gliding locomotion in vertebrates (Dial et al. 2004).

Forest structure has long been recognized as a key factor shaping the distribution of gliding vertebrates. At regional scales, studies on specific taxa have shown that canopy physiognomy, such as canopy height and coverage, can constrain species distributions even in climatically suitable areas (Ishwar et al. 2003; Ramamoorthi and Shai 2021; Abedin et al. 2025a, b). Large-scale analyses have provided empirical support for this idea, demonstrating that gliding vertebrate richness is positively associated with tree height and influenced by forest structural attributes (Emmons and Gentry 1983; Dial et al. 2004; Corlett 2007; Wagner et al. 2023).

However, these studies remain taxonomically or spatially limited, often focusing on a single clade or restricted geographic regions, which limits our ability to identify general patterns across vertebrate groups. As gliding has evolved independently across multiple lineages, examining these taxa collectively provides an opportunity to assess whether common environmental constraints underlie this locomotion strategy.

Understanding these constraints is particularly important in Southeast Asia, where rapid deforestation and land-use change are transforming forest ecosystems at an unprecedented rate. The region has experienced some of the highest levels of forest loss globally, with severe consequences for biodiversity (Sodhi et al. 2004; Ashton and Kettle 2012). Because gliding locomotion enhances predator avoidance, foraging efficiency, and mating success (Dial 2003), gliding vertebrates are expected to be especially sensitive to such changes as their locomotion depends on the three-dimensional structure and connectivity of forest canopies (Suzuki and Yanagawa 2019; Suzuki 2023). Habitat degradation, fragmentation, and changes in canopy structure may therefore have disproportionate impacts on their distributions.

In addition to habitat loss, climate change is altering temperature and precipitation regimes across tropical regions (Intergovernmental Panel on Climate Change (IPCC) 2023; Ombadi et al. 2026), further influencing species distributions. While climate has been widely incorporated into species distribution models, forest structural variables have often been examined separately, and their combined effects remain poorly understood, particularly when evaluated across multiple taxa.

In this study, we investigate the environmental determinants of gliding vertebrate distributions in Southeast Asia using species occurrence data and environmental variables, including climate and forest structure. By integrating multiple vertebrate groups within a single analytical framework, we aim to identify both shared and taxon-specific environmental drivers and to evaluate whether consistent environmental constraints emerge across independently evolved gliding lineages.

## Methods

### Occurrence data collection and processing

Occurrence records of gliding vertebrates in Southeast Asia were obtained from the Global Biodiversity Information Facility (GBIF) (GBIF.org 2026). Target taxa included flying lemurs, flying squirrels, gliding lizards, gliding snakes, and gliding frogs. Occurrence data were acquired using genus-level queries, with each download conducted separately. Because occurrence records were unevenly distributed across species, analyses were conducted at the genus level to ensure sufficient sample sizes and to reduce bias caused by uneven sampling among species. The full list of genera used in this study is shown in Table S1.

The study area was restricted to 90–135°E and 15°S–35°N to focus on Southeast Asia and exclude northern China. Only records with geographic coordinates and no known geospatial issues were retained, and records prior to 2000 were excluded to match the temporal coverage of environmental data. Several genera (e.g., *Aeretes, Biswamoyopterus, Eoglaucomys, Eupetaurus, Petinomys, Pteromys, Trogopterus*, and *Ptychozoon*) were excluded due to the absence of occurrence records, which may reflect data limitations rather than true absence.

Data cleaning was conducted in multiple steps. Records with coordinate uncertainty greater than 10 km were removed. Additionally, records located at country centroids, capitals, biodiversity institutions, equal latitude-longitude coordinates, zero coordinates, and marine locations were excluded. To reduce spatial autocorrelation, occurrence points were thinned to a minimum distance of 10 km for each genus using projected coordinates, which improves independence among occurrence points. The cleaned data are shown in Tables 1 and 2. This resulted in a reduction in sample size, but improved spatial independence among records.

**Table 1.** Summary of occurrence records. Number of occurrence records for each genus before cleaning, after cleaning, and after spatial thinning (10 km minimum distance). Genera are grouped into five categories: Flying lemurs, Flying squirrels, Gliding lizards, Gliding snakes, and Gliding frogs.

| Genus | Group | N before cleaning | N after cleaning | N after thinning |
| --- | --- | --- | --- | --- |
| Cynocephalus | Flying lemurs | 100 | 98 | 7 |
| Galeopterus | Flying lemurs | 1495 | 1118 | 67 |
| Aeromys | Flying squirrels | 247 | 244 | 16 |
| Hylopetes | Flying squirrels | 40 | 35 | 25 |
| Iomys | Flying squirrels | 18 | 17 | 6 |
| Petaurillus | Flying squirrels | 4 | 4 | 2 |
| Petaurista | Flying squirrels | 1051 | 962 | 151 |
| Petinomys | Flying squirrels | 38 | 38 | 4 |
| Pteromyscus | Flying squirrels | 1 | 1 | 1 |
| Draco | Gliding lizards | 4238 | 3496 | 637 |
| Chrysopelea | Gliding snakes | 1960 | 1218 | 419 |
| Rhacophorus | Gliding frogs | 1592 | 1581 | 241 |

**Table 2.** Summary of occurrence records by groups. Total number of occurrence records for each group before cleaning, after cleaning, and after spatial thinning (10 km minimum distance).

| Group | N before cleaning | N after cleaning | N after thinning |
| --- | --- | --- | --- |
| Flying lemurs | 1595 | 1216 | 74 |
| Flying squirrels | 1399 | 1301 | 205 |
| Gliding lizards | 4238 | 3496 | 637 |
| Gliding snakes | 1960 | 1218 | 419 |
| Gliding frogs | 1592 | 1581 | 241 |

### Environmental variables

Environmental predictors included 19 bioclimatic variables obtained from WorldClim version 2.1 (Fick and Hijmans 2017) and elevation obtained from Global multi-resolution terrain elevation data 2010 (GMTED2010) (Jeffrey and Dean 2011). Forest cover was derived from the Hansen Global Forest Change dataset (treecover2000) (Hansen et al. 2013). Canopy structure was further characterized using GEDI Level 3 data, including mean canopy height and its standard deviation (Dubayah et al. 2021). The descriptions and native resolutions of each variable are shown in Table S2. Because environmental datasets were derived from different time periods (e.g., WorldClim representing averages for 1970–2000, treecover2000 for the year 2000, and GEDI data from 2019 onward), occurrence records were restricted to those collected after 2000 to minimize temporal mismatch between occurrence and environmental data. WorldClim variables represent long-term climatic averages and are therefore widely used for species distribution modeling despite temporal differences (Fick and Hijmans 2017).

Environmental values were extracted for each occurrence point by calculating the mean of raster values within a 5 km buffer. The buffer size was selected to match the spatial resolution of environmental variables and to capture local environmental conditions surrounding each occurrence record, while accounting for potential spatial uncertainty in occurrence data. Records with missing environmental values were excluded from subsequent analyses.

### Statistical analyses

All statistical analyses were conducted using R version 4.4.1 (R Core Team 2024) with packages car (Fox and Weisberg 2019), caret (Kuhn 2008), CoordinateCleaner (Zizka et al. 2019), geodata (Hijmans 2025), rgbif (Chamberlain and Boettiger 2017; Chamberlain et al. 2026), and terra (Hijmans 2026). To evaluate environmental associations, presence–background logistic regression models were constructed for each group. In these models, occurrence records were treated as presence data, whereas background points represented available environmental conditions within the study region rather than true absences. Therefore, model coefficients should be interpreted as relative associations with environmental variables, rather than absolute probabilities of occurrence.

A total of 10,000 background points were sampled uniformly within the bounding box (90–135°E, 15°S–35°N). The same number of background points (n = 10,000) was used for all group-specific models. This number was chosen to adequately represent environmental variation across the study region. Environmental variables for background points were extracted using the same procedure as for occurrence records by calculating the mean values within a 5 km buffer around each point. Background points with missing environmental values were excluded to ensure consistency with occurrence data. This approach assumes that background points approximate the environmental availability within the study region, although this assumption may be affected by sampling bias in occurrence data.

A baseline model (m1) was constructed using a subset of ecologically representative variables selected to represent major climatic gradients, including temperature and precipitation-related variables. A second model (m2) was constructed by adding elevation to the m1 model to account for topographic effects. Multiple candidate models (m3–m6) were defined based on different thresholds of pairwise Pearson’s correlation coefficients (|r| < 0.7, 0.75, 0.8, and 0.9). The full composition of each candidate model is provided in Table S3. Model performance was compared using the Akaike information criterion (AIC) (Akaike 1974), and multicollinearity was assessed using variance inflation factors (VIFs).

## Results

### Distribution patterns

The spatial distribution of occurrence records across taxa is shown in Figure 1. Flying lemurs were primarily recorded in the Philippines and Malaysia. Flying squirrels were most frequently recorded in Taiwan. Gliding lizards and snakes were widely recorded across tropical Southeast Asia. Most gliding snake records were concentrated at lower latitudes (approximately 10°S to 10°N). Gliding frogs were mainly recorded within the Indochinese peninsula.

**Figure 1.**
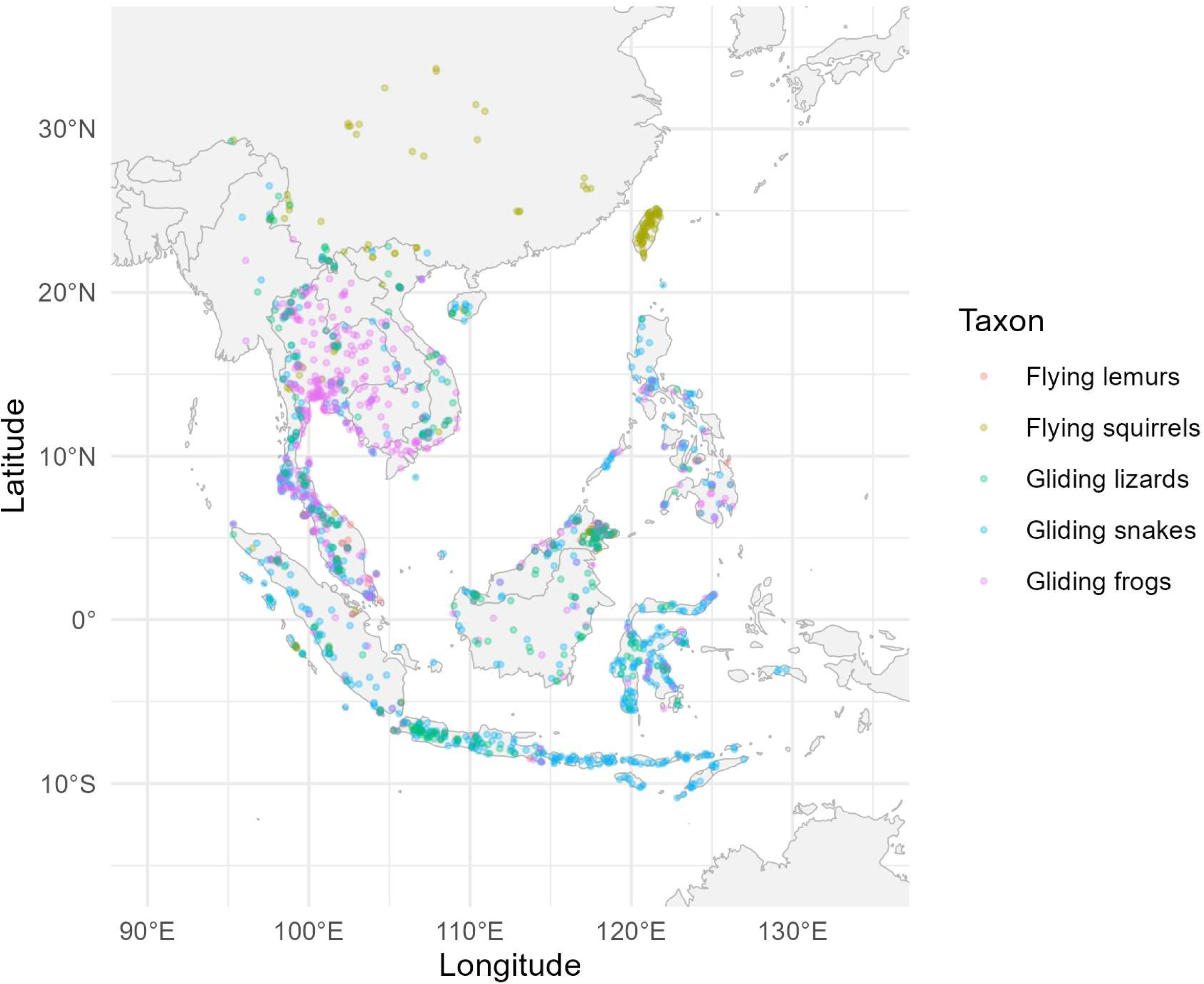
Geographic distribution of gliding vertebrates in Southeast Asia. Points represent cleaned GBIF occurrence data for five groups: Flying lemurs (red), Flying squirrels (yellow), Gliding lizards (green), Gliding snakes (blue), and Gliding frogs (purple).

### Model performance

Model comparisons based on AIC indicated that more complex models generally provided the best fit across taxonomic groups (Table S4). However, these models exhibited substantial multicollinearity among predictors, as indicated by high VIF values (Table S4).

Although VIF thresholds are context-dependent and should not be used as strict criteria (O’Brien 2007; Morrissey and Ruxton 2018; Marcoulides and Raykov 2019), models with lower multicollinearity were preferred to ensure more stable parameter estimation and interpretability. The composition of each model is shown in Table S3.

While model m3 did not achieve the lowest AIC, it consistently showed substantially lower multicollinearity and a reduced number of predictors. Given that the primary aim of this study is to identify and interpret environmental drivers rather than to maximize predictive performance, we prioritized model interpretability and the stability of parameter estimates.

Therefore, model m3 was selected for further interpretation as a compromise between model fit and reduced multicollinearity. Simpler models such as m1 showed low multicollinearity but substantially poorer model fit, and were therefore not considered suitable.

### Environmental drivers

Figure 2 illustrates the estimated effects of environmental variables in model m3. Across taxa, temperature seasonality showed consistently negative effects, while mean canopy height showed positive effects.

**Figure 2.**
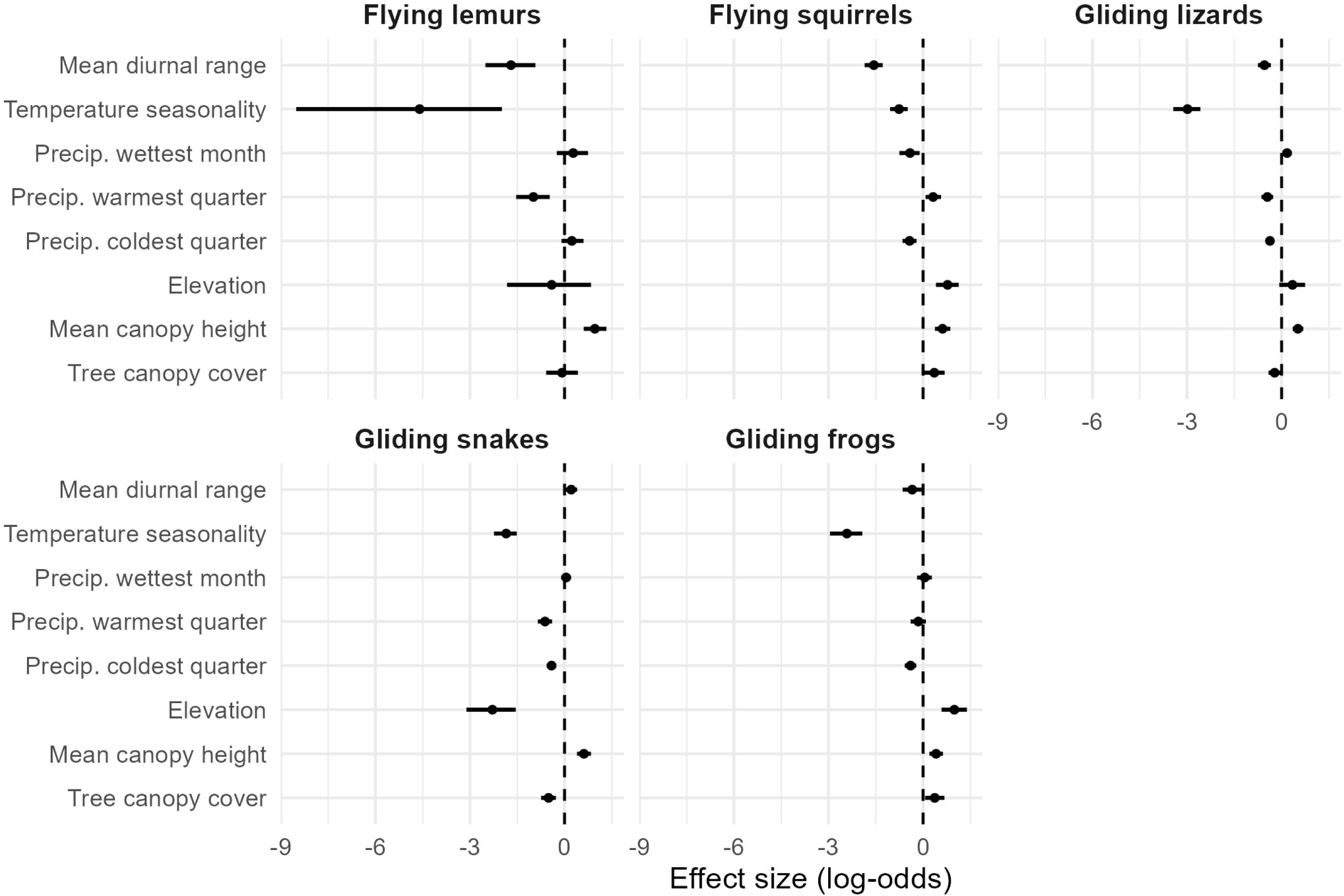
Environmental effects on occurrence based on model m3. Estimated effects of environmental variables on the occurrence of gliding vertebrates based on the logistic regression model m3. Points indicate estimated regression coefficients, and horizontal lines represent 95% confidence intervals. The dashed vertical line indicates zero effect. In the variable labels, “Temp.” and “Precip.” denote temperature and precipitation respectively.

Environmental effects also differed in both magnitude and direction among taxa. Flying lemurs showed a relatively strong negative effect of temperature seasonality and a positive effect of canopy height, while other variables showed relatively weak effects. Flying squirrels showed positive effects of elevation, canopy height, and canopy cover, and negative effects of mean diurnal range and temperature seasonality. Gliding lizards showed a negative effect of temperature seasonality, with generally minor effects of other variables. Gliding snakes showed negative effects of temperature seasonality and elevation, whereas other variables had weak effects. Gliding frogs showed a negative effect of temperature seasonality and positive effects of elevation and canopy height.

## Discussion

In this study, we examined the distribution patterns and environmental drivers of gliding vertebrates in Southeast Asia. Our results reveal both shared and taxon-specific environmental responses, indicating that although gliding has evolved convergently across vertebrates, its ecological associations are divergent.

## Distribution patterns

The spatial distribution of occurrence records generally reflected known biogeographic patterns but also highlighted potential biases in data availability. Flying lemurs were primarily recorded in the Philippines and Malaysia (Figure 1), consistent with their known distribution comprising *Cynocephalus volans*, which is endemic to the southern Philippines, and *Galeopterus variegatus*, which is distributed across mainland Southeast Asia (Nowak 1999).

In contrast, flying squirrel records were disproportionately concentrated in Taiwan (Figure 1), despite their broad distribution across Southeast Asia (Nowak 1999). This likely reflects sampling and observation biases rather than true distribution patterns. Nocturnal behavior (Nowak 1999), regional accessibility, and uneven survey effort may contribute to this pattern, while several genera known from Southeast Asia were not represented in the dataset (see Methods section), further suggesting data limitations.

Gliding lizards showed a broad distribution across both mainland and insular Southeast Asia (Figure 1), consistent with the widespread distribution of the genus *Draco* (McGuire and Dudley 2011). However, their detectability may be reduced due to camouflage of the patagium (Klomp et al. 2014) and potentially low population densities in some species (IUCN 2025). Gliding snakes also exhibited a wide distribution but were more frequently recorded at lower latitudes (Figure 1), suggesting a stronger association with equatorial tropical environments.

Gliding frogs were primarily recorded in mainland Southeast Asia, particularly in the Indochinese region (Figure 1). However, amphibians are often underrepresented in large-scale biodiversity databases, and this pattern may partly reflect limited survey effort and the high proportion of undescribed or poorly documented species in the region, as well as ongoing declines driven by habitat loss and other threats (Rowley et al. 2009).

## Environmental drivers

To better understand the ecological meaning of these patterns, we interpret environmental effects in light of functional and ecological differences among taxa. A consistent pattern across all groups was the negative effect of temperature seasonality and the positive effect of canopy height (Figure 2). These results indicate that gliding vertebrates preferentially occur in climatically stable environments with well-developed vertical forest structure. This pattern is consistent with the predominance of aseasonal tropical climates in Southeast Asia and the functional requirements of gliding locomotion, which depend on sufficient height for effective movement and landing (Suzuki 2023). In addition, high forest canopy height facilitates three-dimensional movement and increases resource availability in arboreal animals, highlighting the importance of vertical forest structure for gliding vertebrates (Feng et al. 2020; Harel et al. 2022; Yang et al. 2023).

In contrast, tree canopy cover showed little or no significant effect among taxa. Although canopy closure is often assumed to facilitate arboreal connectivity (Ishwar et al. 2003; Davies et al. 2017; Ramamoorthi and Shai 2021; Harel et al. 2022; Wagner et al. 2023), our results suggest that vertical forest structure is more important than horizontal canopy continuity. This distinction highlights that not all aspects of forest structure are equally relevant to gliding animals, suggesting that vertical space may be more critical than horizontal connectivity for gliding locomotion.

The relatively uniform response to temperature seasonality may also reflect limited environmental variation within the study region (Janzen 1967), potentially constraining its explanatory power.

## Taxon-specific responses

Despite shared associations with climatic stability and forest structure, environmental responses differed among taxa.

In flying lemurs, most variables showed weak or non-significant effects (Figure 2), likely due in part to limited sample size. In flying squirrels, most variables were statistically significant but had relatively small effect sizes (Figure 2). Because a large portion of records belonged to the genus *Petaurista* (Table 1), these results may primarily reflect the ecological characteristics of this lineage. Notably, precipitation of the wettest month showed a negative effect only in this group, although the underlying mechanism remains unclear. It may reflect avoidance of extremely high rainfall conditions, which could interfere with gliding performance caused by wetting of fur, a characteristic unique to mammals, or alter resource availability during peak rainfall periods. Also, it may reflect an indirect association with other environmental factors correlated with precipitation, rather than a direct causal effect.

Gliding lizards exhibited weak associations with most environmental variables, including elevation (Figure 2). Although they are often reported to occur below certain elevation thresholds (Ishwar et al. 2003; Venugopal 2010), this pattern was not strongly reflected in the model. This may be because high-elevation environments (over 2,000m) are relatively rare within the study region, and *Draco* is largely restricted to lower elevations (Ishwar et al. 2003; Venugopal 2010), reducing variation along the elevation gradient and weakening its effects in the model.

In gliding snakes, elevation showed a clear negative effect (Figure 2), indicating a preference for lowland environments. This pattern, which was not observed in other taxa, suggests a tendency toward lowland tropical forests. As ectotherms, snakes are influenced by ambient temperature, and cooler conditions at higher elevations may limit activity, locomotor performance, or foraging opportunities (Angilletta 2009). In addition, because *Chrysopelea* are carnivorous and prey on small mammals or birds, their occurrence may also be indirectly constrained by the elevational distributions of prey.

For gliding frogs, precipitation variables did not show stronger effects than in other groups, despite amphibians being highly dependent on moisture conditions. This discrepancy likely reflects a mismatch between macroclimatic precipitation and microhabitat conditions critical for amphibians. Precipitation does not necessarily reflect local humidity and water availability, and thus may not fully capture the moisture environment. Additionally, because most of Southeast Asia experiences high rainfall, limited variation in precipitation may reduce its explanatory power.

## Synthesis

Overall, our results indicate that gliding vertebrates share a common association with climatically stable tropical environments, but their responses to other environmental variables differ substantially among taxa. This highlights that convergence in locomotor function does not necessarily imply convergence in ecological niches. Instead, environmental associations are shaped by a combination of phylogenetic constraints, locomotor function, ecological traits, and life history differences, highlighting the complexity of ecological constraints underlying convergent locomotor strategies.

These findings may also have implications for conservation, as taxa with more specialized environmental associations, such as gliding snakes, may be particularly vulnerable to habitat alteration.

## Acknowledgements

We gratefully acknowledge that this study was conceptualized through the network and community built within the Nippon Foundation HUMAI Program, which provided a valuable intellectual environment. We thank Takumi Taga for sharing valuable comments on the manuscript.

## Conflict of interest statement

The authors have no conflicts of interest to disclose.

## Data availability statement

All scripts and processed data used for the analyses are available on GitHub and archived via Zenodo (DOI: 10.5281/zenodo.19912184).

## Supporting information

**Table S1.** List of GBIF data sources. List of genera, corresponding GBIF taxon keys, and download DOIs for occurrence data used in the analyses.

**Table S2.** Details of environmental variables. List of the environmental variables used in the analyses, including bioclimatic variables, elevation, and forest structure variables, along with their descriptions, data sources, and native spatial resolutions.

**Table S3.** Candidate models and their predictor variables. List of the candidate logistic regression models and the environmental variables included in each model used for model selection.

**Table S4.** Model comparison based on Akaike information criterion (AIC) and multicollinearity diagnostics. Comparison of the candidate logistic regression models for each group using AIC and variance inflation factors (VIFs). Delta AIC values are calculated relative to the model with the lowest AIC within each group. For each model, the maximum VIF among predictor variables is shown. Model m3 was selected for further interpretation as it showed low multicollinearity while maintaining relatively low AIC values.

## Notes

https://doi.org/10.5281/zenodo.19912184

### Competing Interest Statement

The authors have declared no competing interest.

