## supplementary tables for "Convergent gliding, divergent ecology: Environmental drivers of gliding vertebrates in Southeast Asia"

Table S1

| Genus | GBIF taxon key | Download DOI |
|---|---|---|
| Cynocephalus | 2432089 | https://doi.org/10.15468/dl.tjv5c3 |
| Galeopterus | 2432086 | https://doi.org/10.15468/dl.5ujszf |
| Aeromys | 2437382 | https://doi.org/10.15468/dl.7z7d3w |
| Hylopetes | 2437527 | https://doi.org/10.15468/dl.3wpu3n |
| Iomys | 2437355 | https://doi.org/10.15468/dl.bfrh42 |
| Petaurillus | 2437558 | https://doi.org/10.15468/dl.wsp4j8 |
| Petaurista | 2437549 | https://doi.org/10.15468/dl.en3yz6 |
| Petinomys | 2437248 | https://doi.org/10.15468/dl.nvkfaj |
| Pteromyscus | 2437226 | https://doi.org/10.15468/dl.5rct6r |
| Draco | 2466605 | https://doi.org/10.15468/dl.ydt7cp |
| Chrysopelea | 2457099 | https://doi.org/10.15468/dl.axp7md |
| Rhacophorus | 2421577 | https://doi.org/10.15468/dl.jmt8pw |

Table S2

| Code | Description | Source | Native Resolution |
|---|---|---|---|
| BIO1 | Annual Mean Temperature | WorldClim v2.1 | 2.5 arc-min |
| BIO2 | Mean Diurnal Range (Mean of monthly (max temp - min temp)) | WorldClim v2.1 | 2.5 arc-min |
| BIO3 | Isothermality (BIO2 / BIO7) (×100) | WorldClim v2.1 | 2.5 arc-min |
| BIO4 | Temperature Seasonality (standard deviation ×100) | WorldClim v2.1 | 2.5 arc-min |
| BIO5 | Max Temperature of Walmest Month | WorldClim v2.1 | 2.5 arc-min |
| BIO6 | Min Temperature of Coldest Month | WorldClim v2.1 | 2.5 arc-min |
| BIO7 | Temperature Annual Range (BIO5 - BIO6) | WorldClim v2.1 | 2.5 arc-min |
| BIO8 | Mean Temperature of Wettest Quarter | WorldClim v2.1 | 2.5 arc-min |
| BIO9 | Mean Temperature of Driest Quarter | WorldClim v2.1 | 2.5 arc-min |
| BIO10 | Mean Temperature of Warmest Quarter | WorldClim v2.1 | 2.5 arc-min |
| BIO11 | Mean Temperature of Coldest Quarter | WorldClim v2.1 | 2.5 arc-min |
| BIO12 | Annual Precipitation | WorldClim v2.1 | 2.5 arc-min |
| BIO13 | Precipitation of Wettest Month | WorldClim v2.1 | 2.5 arc-min |
| BIO14 | Precipitation of Driest Month | WorldClim v2.1 | 2.5 arc-min |
| BIO15 | Precipitation Seasonality (Coefficient of Variation) | WorldClim v2.1 | 2.5 arc-min |
| BIO16 | Precipitation of Wettest Quarter | WorldClim v2.1 | 2.5 arc-min |
| BIO17 | Precipitation of Driest Quarter | WorldClim v2.1 | 2.5 arc-min |
| BIO18 | Precipitation of Warmest Quarter | WorldClim v2.1 | 2.5 arc-min |
| BIO19 | Previpitation of Coldest Quarter | WorldClim v2.1 | 2.5 arc-min |
| elev | Elevation | Global Multi-resolution Terrain Elevation Data 2010 | 2.5 arc-min |
| forest_cover | Tree Canopy Cover | Hansen Global Forest Change v1.12 | 30m |
| ch_mean | Mean Canopy Height | GEDI L3 Gridded Land Surface Metrics v2 | 1km |
| ch_sd | Standard Deviation of Canopy Height | GEDI L3 Gridded Land Surface Metrics v2 | 1km |

Table S3

| Model | Variables |
|---|---|
| m1 | BIO1, BIO4, BIO12, BIO15, forest_cover |
| m2 | BIO1, BIO4, BIO12, BIO15, elev, forest_cover |
| m3 | BIO2, BIO4, BIO13, BIO18, BIO19, elev, forest_cover, ch_mean |
| m4 | BIO2, BIO4, BIO13, BIO15, BIO18, BIO19, elev, ch_sd |
| m5 | BIO2, BIO4, BIO12, BIO13, BIO15, BIO18, BIO19, elev, forest_cover, ch_sd |
| m6 | BIO2, BIO4, BIO7, BIO12, BIO14, BIO15, BIO16, BIO18, BIO19, elev, forest_cover, ch_mean, ch_sd |

Table S4

| Model | df | AIC | Delta AIC | max VIF |
|---|---|---|---|---|
| **Flying lemurs** | | | | |
| m1 | 6 | 546.2654 | 85.2974 | 2.2128 |
| m2 | 7 | 533.3814 | 72.4134 | 11.1361 |
| m3 | 9 | 481.9418 | 20.9738 | 3.1345 |
| m4 | 10 | 463.8053 | 2.8373 | 5.4264 |
| m5 | 11 | 460.9680 | 0.0000 | 12.8918 |
| m6 | 14 | 466.5093 | 5.5413 | 25.9696 |
| **Flying squirrels** | | | | |
| m1 | 6 | 1422.3094 | 205.8881 | 2.4705 |
| m2 | 7 | 1401.3516 | 184.9302 | 48.6242 |
| m3 | 9 | 1252.4468 | 36.0255 | 3.7926 |
| m4 | 10 | 1244.8500 | 28.4286 | 6.9344 |
| m5 | 11 | 1244.7348 | 28.3134 | 15.4369 |
| m6 | 14 | 1216.4214 | 0.0000 | 121.2093 |
| **Gliding lizards** | | | | |
| m1 | 6 | 2467.7344 | 111.3823 | 2.4730 |
| m2 | 7 | 2453.2198 | 96.8677 | 16.3717 |
| m3 | 9 | 2455.3813 | 99.0292 | 3.4091 |
| m4 | 10 | 2362.3215 | 5.9694 | 6.5484 |
| m5 | 11 | 2363.6526 | 7.3005 | 12.4015 |
| m6 | 14 | 2356.3521 | 0.0000 | 50.1551 |
| **Gliding snakes** | | | | |
| m1 | 6 | 1901.6466 | 16.1341 | 2.8966 |
| m2 | 7 | 1885.5125 | 0.0000 | 9.9343 |
| m3 | 9 | 1942.1469 | 56.6344 | 4.3324 |
| m4 | 10 | 1902.1931 | 16.6806 | 6.2577 |
| m5 | 11 | 1886.3841 | 0.8716 | 14.3388 |
| m6 | 14 | 1892.7655 | 7.2530 | 60.6934 |
| **Gliding frogs** | | | | |
| m1 | 6 | 1317.0910 | 17.8733 | 3.8044 |
| m2 | 7 | 1316.0633 | 16.8457 | 33.0341 |
| m3 | 9 | 1309.1356 | 9.9180 | 2.9220 |
| m4 | 10 | 1300.2025 | 0.9849 | 8.1279 |
| m5 | 11 | 1301.3134 | 2.0958 | 13.6326 |
| m6 | 14 | 1299.2176 | 0.0000 | 67.0704 |
